## Supplemental Tables 1-4 for "Feasibility of Integrating Canine Olfaction with Chemical and Microbial Profiling of Urine to Detect Lethal Prostate Cancer"

**Table S1. Sample use among canine olfaction, GC-MS/ANN, and microbial profiling.**

| **Sample ID** | **Age** | **PSA** | **Status** | **Dog Training** | **Dog Trial** | **GC-MS/ ANN** | **Microbiome** |
| --- | --- | --- | --- | --- | --- | --- | --- |
| AWP-5568 | 60 | 3.3 | Control |  |  |  |  |
| AWP-5577 | 45 | 1.6 | Control |  |  |  |  |
| AWP-5585 | 51 | 10 | Control |  |  |  |  |
| AWP-5648 | 59 | 3.56 | Control |  |  |  |  |
| AWP-5679 | 51 | 1.1 | Control |  |  |  |  |
| AWP-5726 | 53 | 6.5 | Control |  |  |  |  |
| AWP-5742 | 79 | 6 | Control |  |  |  |  |
| AWP-5767 | 78 | 2.86 | Control |  |  |  |  |
| AWP-5936 | 64 | 17 | Control |  |  |  |  |
| AWP-5937 | 51 | 12 | Control |  |  |  |  |
| AWP-5986 | 69 | 10.8 | Control |  |  |  |  |
| AWP-6147 | 57 | 4.9 | Control |  |  |  |  |
| AWP-6195 | 55 | 1.2 | Control |  |  |  |  |
| AWP-6201 | 70 | 1.9 | Control |  |  |  |  |
| AWP-6241 | 52 | 10 | Control |  |  |  |  |
| AWP-6255 | 56 | 4.48 | Control |  |  |  |  |
| AWP-6258 | 57 | 4.8 | Control |  |  |  |  |
| AWP-6598 | 80 | 14.2 | Control |  |  |  |  |
| AWP-6651 | 67 | 7.5 | Control |  |  |  |  |
| AWP-8135 | 63 | 3.9 | Control |  |  |  |  |
| AWP-8426 | 63 | 3.4 | Control |  |  |  |  |
| AWP-8926 | 51 | 2.9 | Control |  |  |  |  |
| AWP-9211 | 63 | 4.2 | Control |  |  |  |  |
| JHBUI-1100 | 65 | 4.5 | Control |  |  |  |  |
| JHBUI-1980 | 56 | 3.9 | Control |  |  |  |  |
| JHBUI-2181 | 53 | 3.5 | Control |  |  |  |  |
| JHBUI-2510 | 60 | 3.4 | Control |  |  |  |  |
| JHBUI-2980 | 56 | 8.1 | Control |  |  |  |  |
| JHBUI-3039 | 54 | 6.4 | Control |  |  |  |  |
| JHBUI-3061 | 50 | 4.36 | Control |  |  |  |  |
| JHBUI-3179 | 62 | 4.7 | Control |  |  |  |  |
| JHBUI-3255 | 55 | 18.4 | Control |  |  |  |  |
| JHBUI-3381 | 58 | 14.7 | Control |  |  |  |  |
| JHBUI-3420 | 62 | 5.1 | Control |  |  |  |  |
| JHBUI-3422 | 67 | 4.68 | Control |  |  |  |  |
| JHBUI-3631 | 56 | 5.5 | Control |  |  |  |  |
| JHBUI-620 | 73 | Unknown | Control |  |  |  |  |
| JHBUI-735 | 59 | 6.3 | Control |  |  |  |  |
| AWP-5734 | 60 | 5.1 | Cancer |  |  |  |  |
| AWP-6373 | 50 | 4.8 | Cancer |  |  |  |  |
| AWP-9307 | 66 | 7.3 | Cancer |  |  |  |  |
| AWP-9472 | 72 | 8.6 | Cancer |  |  |  |  |
| AWP-9582 | 71 | 3 | Cancer |  |  |  |  |
| JHBUI-1028 | 71 | 53.4 | Cancer |  |  |  |  |
| JHBUI-2175 | 75 | 4.97 | Cancer |  |  |  |  |
| JHBUI-2719 | 73 | 9.27 | Cancer |  |  |  |  |
| JHBUI-2976 | 64 | 76.8 | Cancer |  |  |  |  |
| JHBUI-2978 | 54 | 23.8 | Cancer |  |  |  |  |
| JHBUI-3147 | 49 | 9.2 | Cancer |  |  |  |  |
| JHBUI-887 | 65 | 6.1 | Cancer |  |  |  |  |
|  |  | **Total Controls** |  | **15** | **21** | **30** | **37** |
|  |  | **Total Cancers** |  | **5** | **7** | **6** | **12** |

**Table S2. Summary of canine training data over 22 training days.** **Number and percent correct after each pass.**

| Sample type | Dog | Analysis | Sample ID | Challenges | % correct after 1st pass | % correct after 2nd pass | % correct after 3rd pass | % correct after final pass |
| --- | --- | --- | --- | --- | --- | --- | --- | --- |
| **Biopsy-Negative Controls** | | | | 442 | 83.7% | 76.2% | 76.2% | 76.2% |
|  | **Florin** | |  |  |  |  |  |  |
|  |  | Detection | | 232 | 81.0% | 73.2% | 73.2% | 73.2% |
|  |  |  | AWP-5568 | 27 | 100.0% | 100.0% | 100.0% | 100.0% |
|  |  |  | AWP-5648 | 2 | 50.0% | 50.0% | 50.0% | 50.0% |
|  |  |  | AWP-5742 | 26 | 95.0% | 80.0% | 85.0% | 85.0% |
|  |  |  | AWP-5937 | 13 | 55.6% | 55.6% | 55.6% | 55.6% |
|  |  |  | AWP-6195 | 22 | 68.8% | 56.3% | 50.0% | 50.0% |
|  |  |  | AWP-6255 | 16 | 100.0% | 92.3% | 92.3% | 92.3% |
|  |  |  | AWP-6651 | 13 | 72.7% | 72.7% | 72.7% | 72.7% |
|  |  |  | AWP-9211 | 25 | 64.7% | 35.3% | 35.3% | 35.3% |
|  |  |  | JHBUI-0735 | 26 | 90.5% | 81.0% | 81.0% | 81.0% |
|  |  |  | JHBUI-1980 | 7 | 33.3% | 33.3% | 33.3% | 33.3% |
|  |  |  | JHBUI-2510 | 11 | 100.0% | 100.0% | 100.0% | 100.0% |
|  |  |  | JHBUI-2980 | 23 | 58.8% | 58.8% | 58.8% | 58.8% |
|  |  |  | JHBUI-3039 | 11 | 90.9% | 81.8% | 81.8% | 81.8% |
|  |  |  | JHBUI-3381 | 5 | 100.0% | 100.0% | 100.0% | 100.0% |
|  |  |  | JHBUI-3422 | 5 | 100.0% | 100.0% | 100.0% | 100.0% |
|  | **Midas** | |  |  |  |  |  |  |
|  |  | Detection | | 210 | 86.9% | 79.7% | 79.7% | 79.7% |
|  |  |  | AWP-5568 | 25 | 85.0% | 80.0% | 80.0% | 80.0% |
|  |  |  | AWP-5648 | 5 | 60.0% | 40.0% | 40.0% | 40.0% |
|  |  |  | AWP-5742 | 19 | 84.6% | 84.6% | 84.6% | 84.6% |
|  |  |  | AWP-5937 | 12 | 85.7% | 85.7% | 85.7% | 85.7% |
|  |  |  | AWP-6195 | 13 | 87.5% | 87.5% | 87.5% | 87.5% |
|  |  |  | AWP-6255 | 13 | 100.0% | 87.5% | 87.5% | 87.5% |
|  |  |  | AWP-6651 | 13 | 80.0% | 80.0% | 80.0% | 80.0% |
|  |  |  | AWP-9211 | 20 | 84.6% | 76.9% | 76.9% | 76.9% |
|  |  |  | JHBUI-0735 | 20 | 94.4% | 88.9% | 88.9% | 88.9% |
|  |  |  | JHBUI-1980 | 9 | 100.0% | 100.0% | 100.0% | 100.0% |
|  |  |  | JHBUI-2510 | 10 | 100.0% | 85.7% | 85.7% | 85.7% |
|  |  |  | JHBUI-2980 | 23 | 84.2% | 68.4% | 68.4% | 68.4% |
|  |  |  | JHBUI-3039 | 10 | 100.0% | 80.0% | 80.0% | 80.0% |
|  |  |  | JHBUI-3381 | 9 | 87.5% | 87.5% | 87.5% | 87.5% |
|  |  |  | JHBUI-3422 | 9 | 50.0% | 50.0% | 50.0% | 50.0% |
| **Cancer** | |  |  | 152 | 49.6% | 61.7% | 64.5% | 65.2% |
|  | **Florin** | |  |  |  |  |  |  |
|  |  | Detection | | 80 | 47.9% | 59.2% | 60.6% | 62.0% |
|  |  |  | AWP-5734 | 6 | 50.0% | 66.7% | 66.7% | 66.7% |
|  |  |  | AWP-6373 | 13 | 72.7% | 72.7% | 72.7% | 72.7% |
|  |  |  | AWP-9472 | 24 | 47.6% | 71.4% | 76.2% | 81.0% |
|  |  |  | JHBUI-0887 | 25 | 47.8% | 47.8% | 47.8% | 47.8% |
|  |  |  | JHBUI-2978 | 12 | 20.0% | 40.0% | 40.0% | 40.0% |
|  | **Midas** | |  |  |  |  |  |  |
|  |  | Detection | | 72 | 51.4% | 64.3% | 68.6% | 68.6% |
|  |  |  | AWP-5734 | 7 | 57.1% | 57.1% | 57.1% | 57.1% |
|  |  |  | AWP-6373 | 14 | 50.0% | 64.3% | 64.3% | 64.3% |
|  |  |  | AWP-9472 | 18 | 77.8% | 88.9% | 88.9% | 88.9% |
|  |  |  | JHBUI-0887 | 22 | 30.0% | 45.0% | 55.0% | 55.0% |
|  |  |  | JHBUI-2978 | 11 | 45.5% | 63.6% | 72.7% | 72.7% |
| **Grand Total** |  |  |  | **594** | **73.6%** | **71.9%** | **72.7%** | **72.9%** |

Detection = Indication recorded as a true positive or a false positive and in definition, an operant behavior shown when the dog believes that the odor being sniffed is the correct odor to respond to.

The table shows all training data from 10/24/2018-12/17/2018, with a total number of 22 training days. The data outlines the number of exposures (sniffs) each sample received over the training period, along with sensitivity and specificity from combined exposures after each pass. The number of exposures fluctuates with each sample, with some presented to the dogs significantly less than others. This is partly due to when the samples were unblinded to the team for use in training (samples that were unblinded earlier generally had more exposures), but in some instances an issue with the individual sample may have influenced the decision to remove a sample from a line.

Sensitivity and specificity for each sample may have also been influenced by the quality of the aliquot. Due to the limited availability of samples for training, some aliquots were reused up to 3 times which we believe may have caused a deterioration of learned target stimulus.

**Table S3. Instrument parameters for HS GC-MS testing of urine samples.**

| **Headspace Conditions** | |
| --- | --- |
| Temperature Zones | |
| Oven Temperature (°C) | 90 |
| Loop Temperature (°C) | 110 |
| Tr. Line Temperature (°C) | 130 |
| Timing | |
| GC Cycle Time (min) | 42 |
| Vial Eq. Time (min) | 60.00 |
| Pressurization Time (min) | 0.15 |
| Loop Fill Time (min) | 0.15 |
| Loop Eq. Time (min) | 0.05 |
| Inject Time (min) | 1.00 |
| Other | |
| Vial Pressure (psi) | 25 |
| Shake | LOW |
| Vial Size (mL) | 20 |
| **GC Conditions** | |
| Zebron ZB-624 | 30 m x 0.25 mm x 1.4 µm |
| Inlet Temperature (°C) | 250 |
| Septum Pure Flow (mL/min) | 3 |
| Mode | SPLIT |
| Split Ratio | 2:1 |
| Column Flow (mL/min) | 1 |
| Oven Program | Hold 40°C for 3 min, ramp 20°C/min to 240°C |
| Total Run Time (min) | 13 |
| **MS Conditions** | |
| Transfer Line Temperature (°C) | 240 |
| Detection Mode | Scan (m/z 30-350) |

**Table S4. Common contaminant genera in microbiome analyses.**

| Achromobacter | Curtobacterium | Pelomonas |
| --- | --- | --- |
| Acidovorax | Geobacillus | Propionibacterium |
| Agrobacterium | Halomonas | Ralstonia |
| Anoxybacillus | Halorhodospira | Rhodobacter |
| Aquabacter | Herbaspirillum | Shewanella |
| Aquabacterium | Janthinobacterium | Sphingobacterium |
| Aquimonas | Mesorhizobium | Sphingobium |
| Bradyrhizobium | Methylobacillus | Sphingomonas |
| Caulobacter | Methylobacterium | Undibacterium |
| Comamonas | Methylomarinum | Xanthomonas |
| Cryobacterium | Paenibacillus | Diaphorobacter |
| Cupriavidus | Pantoea |  |
