## Supplemental Methods for "Feasibility of Integrating Canine Olfaction with Chemical and Microbial Profiling of Urine to Detect Lethal Prostate Cancer"

**Facilities**

Canine training was performed at the Bio-Detection building 2 at Medical Detection Dogs (MDD), UK. The facilities include kitchen/office with viewing gallery that looks onto a climate-controlled dog training room, climate-controlled sample storage room, sample preparation area and equipment cleaning room. The facility also houses -80 ^o^C, -40 ^o^C and -20 ^o^C sample storage freezers. The sample preparation and cleaning room contains a work bench for sample preparation and equipment to facilitate sample processing and cleaning, including a standard domestic refrigerator, a dedicated sink, an ultrasonic cleaner, a commercial dishwasher and an autoclave.

Within the training room there is close circuit television (CCTV) linked to a DVD recorder, a laptop with connected webcam facing the carousel for data recording, and a workspace including a desk. Within the testing arena there is a one- way viewing shield, this allows the project specialist to be concealed during training and testing to avoid the unintended transmission of cues. The shield incorporates one-way glass to allow the project specialist to observe the dog search. This shield also has a flat screen connected to the data collection system. The sample presentation equipment is an eight-position carousel. Due to the limited number of test samples available the pilot study used only four positions in the carousel, positions 1,3,5,7. The carousel and associated pots are surgical grade stainless steel.

After each training and trial session, all equipment holding a sample was thoroughly cleaned via ultrasonic bath filled with enzymatic cleaner (Reprozyme Manual) followed by dishwasher (no detergent) at 80 ºC for three minutes followed by autoclave.

Facility photographs and links to exemplar training and testing videos are provided in the Supplemental Information.

**Canine training protocol**

All training sessions were recorded using an internally developed database system Olfactory Performance Recording Applications (OPRA). Each session was captured on video using a webcam and stored on a protected hard drive at MDD.

Training for the pilot using the supplied JHU samples commenced on November 13, 2018. Initially the training sample set was supplied un-blinded and contained two cancer and six control samples with diagnosis declared. These samples were selected by JHU. At the request of MDD, a second group of three cancer and 9 control training samples were supplied by JHU to aid in the training and calibration of the dogs to these unfamiliar sample types for habituation to the unique sample source. A further change to usual methodology was required to accommodate the limited number of samples available, the usual practice is for samples to be single use only. However, instead of single use the samples were re-used by following a freeze and thaw process for repeated presentation at a later date.

All training data was recorded including video within the database system for future analysis. Due to the limited number of samples available for training, it was decided the training protocol would follow a “forced-choice” scenario with a positive bias for inclusion of a positive sample in each training run. The standard training criteria was adapted to accommodate the limited sample number by removing a control sample if indicated by the dog and replacing it with a blank pot. The dog was then tasked to search again to indicate a further sample. If two wrong choices were made in succession, the run was deemed complete and the training session ended, and the samples were changed.

During the training, samples from the first training group and second training group were selected in order of supply. After sufficient data had been gathered, they were selected by the project specialist’s choice with reference to dogs’ ability to detect or ignore.

On the day of training each sample was selected from the sample list, a labelled glass presentation pot was prepared with details of the sample. The required samples were collected from the freezer and defrosted. The pots and aliquot were placed into the refrigerator to stabilize to an equal temperature of 4 ºC for a minimum of ten minutes before use. Each sample was recovered from the refrigerator, decanted into a glass presentation pot, and sealed with a metal lid. This was left for ten minutes before use. When ready each relevant presentation pot was placed unsealed into the metal pot attached to the carousel in accordance with the requirements for training in that session. All other samples not in use remained sealed until such time that they were required for training. When a sample was removed from the carousel it was resealed with the matched lid until it was required again or was returned to the freezer for future use or storage.

The dogs were trained through various stages of training including, known position and outcome, single-blind and double-blind searches.
